# Bi-parental graph strategy to represent and analyze hybrid plant genomes

**DOI:** 10.1101/2023.11.28.568999

**Authors:** Qianqian Kong, Yi Jiang, Zhiheng Wang, Zijie Wang, Yuting Liu, Yuanxian Gan, Han Liu, Xiang Gao, Xuerong Yang, Xinyuan Song, Hongjun Liu, Junpeng Shi

## Abstract

Hybrid plants are universally existed in wild and often exhibit greater performance of complex traits compared with their parents and other selfing plants. This phenomenon, known as heterosis, has been extensively applied in plant breeding for decades. However, the process of decoding hybrid plant genomes has seriously lagged due to the challenges in their genome assembling and the lack of proper methods to further represent and analyze them. Here we report the assembly and analysis of two hybrids: an intraspecific hybrid between two maize inbred lines and an interspecific hybrid between maize and its wild relative teosinte, based on the combination of PacBio High Fidelity (HiFi) sequencing and chromatin conformation capture sequencing data. The haplotypic assemblies are well-phased at chromosomal scale, successfully resolving the complex loci with extensive parental structural variations (SVs). By integrating into a bi-parental genome graph, the haplotypic assemblies can facilitate downstream short-reads based SV calling and allele-specific gene expression analysis, demonstrating outstanding advantages over one single linear genome. Our work provides an entire workflow which hopefully can promote the deciphering of the large numbers of hybrid plant genomes, especially those whose parents are unknown or unavailable and help to understand genome evolution and heterosis.

## Introduction

So far in wild, the overwhelming majority (>80%) of plant species have remained outcrossed as hybrids, although the transition from hybridization or outcrossing to self-fertilization is believed to be evolutionarily favoured [1, 2]. Hybrids usually exhibit greater performance of complex traits over their parents and other selfing plants, a phenomenon known as heterosis that has been applied in crop breeding programs for several decades [3]. Decoding the genetic basis of heterosis requires allele-wared genome sequences, which in some cases can be acquired by separately assembling and comparing two parental genomes [4-6]. However, in most circumstances, the parents of naturally existing plant hybrids are unknown or unavailable. In the meanwhile, as the only possible option, direct assembling of these heterozygous genomes is still technically challenging by using either short or earlier long noisy reads. Therefore, most genome assemblers adopted a compromised strategy to assemble heterozygous genome by collapsing the sequencing reads from two parental haplotypes into a single “mosaic” set of contigs (Fig. 1), but this strategy ignored parental genetic variants associated with heterosis and allele-specific gene expression [7]. Further advancement has been made to generate the allele-wared genome assemblies, such as in diploid potato [8], tea [9, 10] and lychee [11], but haplotype switch errors were still common for them, especially when assessed at chromosomal or genome-wide scales. In terms of downstream analysis, the previous methods for conventional genome sequences have only represented a single haplotype, indicating the lack of new protocols for these allele-wared or haplotype-resolved assemblies as new genome references.[7]. Although the process of decoding hybrid plant genomes has once far dropped behind, recent innovations of long read sequencing and chromatin conformation capture sequencing technologies (for example, Hi-C), however, enable haplotype-resolved assembly and phasing in hybrids [12, 13], offering a great opportunity to design a new workflow to represent and analyze the large numbers of hybrid plant genomes.

**Fig. 1.**
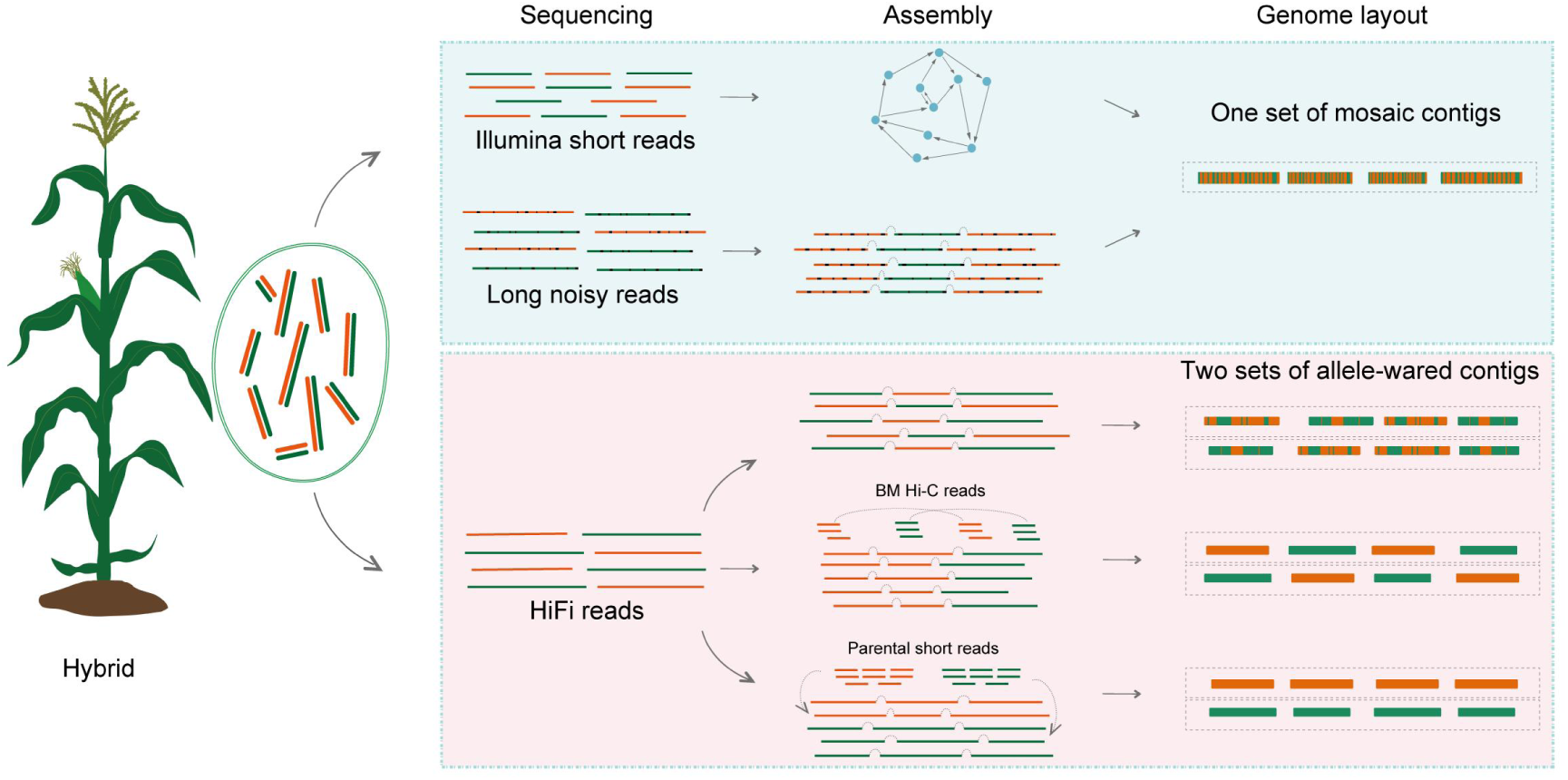
Schematic diagram of genome assembly in plant hybrids. The genome of hybrids comes from the combination of two parental genome, with usually extensive genetic variants. Assembling hybrid genome was technically challenging with short or long noisy reads, therefore most earlier assemblers simply collapsed the reads from two parents into a mosaic set of contigs (**Upper box**). The recent innovation of long read sequencing, especially PacBio HiFi technology, enables haplotype-resolved assemblies by producing two sets of allele-wared contigs (**Bottom box**), although in most cases two haplotypes are switched more-or-less between two set of contigs. Generally, the integration of HiFi long reads with chromatin interaction data (e.g., Hi- C or HiChIP) can generate chromosomal scale, well-phased haplotypic assemblies. However, this strategy may still put paternal and maternal contigs from different chromosomes into a single partition, since Hi-C data was unable to distinguish the paternal and maternal chromosomes in the cells of an offspring. Only when the pedigree information, for example the short reads from two parents were available, can well-phased haplotypic assemblies be further generated at genome-wide scale using the trio-binning strategy.

Here we report high-quality, chromosomal-scale phased assemblies of two plant hybrids with substantially high genome heterozygosity, one from a cross between two typical maizes (*Zea may* ssp. *mays*) inbred lines and the other from an interspecific hybridization between maize and its wild relative teosinte (*Zea may* ssp. *parviglumis*). We provide evidence that these haplotypic assemblies can successfully resolve complex loci with extensive parental structural variations. We finally demonstrate the feasibility and superiority of integrating these haplotypic assemblies into a genome graph, as it can outperform single linear genome in facilitating downstream short-reads based mapping, variant calling and allele-specific gene expression analysis. Hopefully, our work may pave a way to accelerate the process of decoding plant heterozygous genomes and promote their applications as new genome references.

## Results

### Chromosome-scale, well-phased assemblies of highly heterozygous maize genome

A classical maize hybrid that crossed between B73 and Mo17 inbred lines (hereafter denoted as BM) was selected in this study and sequenced by PacBio HiFi technology, considering the reasons as follow: (a) The genome heterozygosity of BM, a key metrics tightly correlated with the difficulty in assembling heterozygous genomes, was estimated to be fairly high (∼2.0%), surpassing most of the published heterozygous genomes in plants, humans and other animals (Table S1). (b) High-quality reference genomes of B73 [14] and Mo17 [4] have been released, which could be used to benchmark our assembly results. We generated a total of ∼118.0 Gb HiFi reads (N50 ∼ 16.1 kb) from 6 SMRT cells, which covered approximately 27x of each haploid maize genome (Table S2 and Fig. S1), according to the experience from maize, human and other representative species [15-17] that approximately 25x to 30x of the HiFi long reads was sufficient for assembling a high-quality reference genome. Subsequently, we tested three popular assemblers, Hifiasm [12], HiCanu [18] and Verkko [19], and found that Hifiasm generated the assemblies with the overall best contiguity, completeness and accuracy (Table S3). Moreover, although all assemblers are able to generate allele-wared assemblies, Hifiasm has developed a built-in function to directly phase the raw assemblies into two sets of primary and alternate contigs, comparing with the assemblies produced by HiCanu or Verkko, which require haplotype purging by additional tools [20]. Due to its better performance, assemblies produced by Hifiasm were chosen in subsequent analysis.

To find out the saturation point of HiFi data on haplotypic assemblies, we sequentially assembled a series of subsampled (10x, 15x, 20x, 25x and all) HiFi reads. The result indicated that the overall size of two haplotypic assemblies converged with the increasing of input data (Fig. S2a), while the assembly quality as assessed by contig N50, BUSCO and LAI were continuously optimized (Fig. S2b-2d), yet still not reaching saturation. Comparing with the latest B73 reference genome and the earlier Mo17 genome assembled by PacBio long noisy reads [4, 14], the final assemblies generated by all HiFi reads exhibited comparable or even better quality (Table 1). As expected, the subregions switches between two haplotypes were still pervasively existed in both hap1 and hap2 assemblies, with particular enrichment in relatively long contigs (Fig. 2a). This phenomenon largely came from mis-joins during contig construction [12], which were technically unavoidable with only HiFi reads as input and may introduce chaos during assembly. To some extent, this may also explain the enlarged size of two haplotypic assemblies (∼2.27 Gb & ∼2.23 Gb) when compared it with the genomes of B73 (∼2.18 Gb) and Mo17 (∼2.18 Gb).

**Fig. 2.**
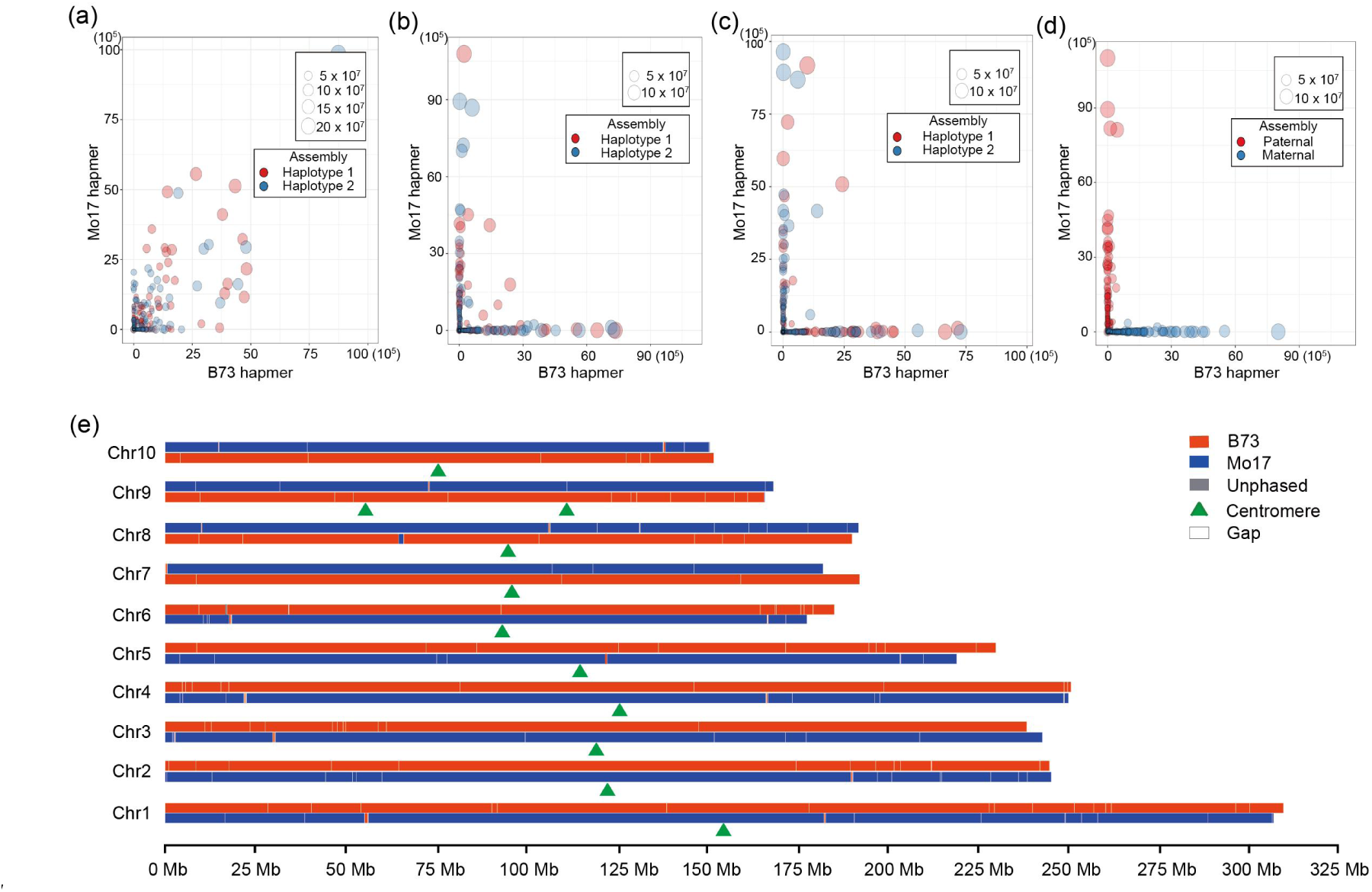
Chromosomal-scale, well-phased assemblies of maize BM genomes. The composition of two haplotypic contigs from B73 and Mo17 was assessed by Merquery for direct assembly **(a), HiFi+HiChIP (b), HiFi+Hi-C (c) and trio-binning (d)**. Each circle represents a contig, and the numbers of paternal-specific and maternal-specific 21-mers were shown in x-axis and y-axis, respectively. **(e)** Chromosome-scale assembly and phasing results for BM assemblies that generated by HiFi+Hi-C method. For each chromosome, the top track and the bottom track indicate hap1 and hap2, respectively.

**Table 1.**
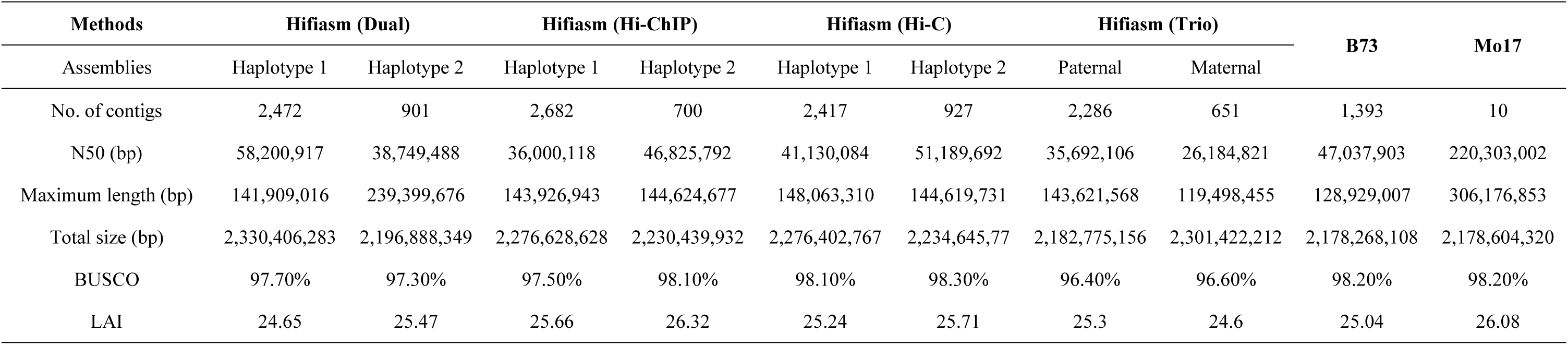
The key features of maize BM genome assemblies.

As the combination of long HiFi reads with chromatin interaction data (for example, Hi-C) has recently been proved to be effective in generating chromosomal-scale, well-phased haplotypic assemblies in human [13], we here attempt to implement it in plant hybrids. Unlike the commonly used trio-binning strategy [21, 22], which utilized the DNA sequences of two parents to initially partition the F1 long reads into two haplotype-specific sets, then assemble each haplotype independently (Fig. 1), the procedure of HiFi+Hi-C can not phase at genome-wide scale (all the chromosomes in one haplotypic assembly came from a single parent), and may still put paternal and maternal contigs from different chromosomes into a single partition, since Hi-C data was still unable to distinguish the paternal and maternal chromosomes in the cells of an offspring [13]. However, this strategy still holds great significance and is probably the best solution at present to decode the haplotype-resolved genomes for individuals without parental information. As a result, we newly generated the Hi-C data (∼96.2 Gb, ∼44x) of maize BM seedlings and collected the HiChIP data (∼69.9 Gb, ∼32x) mediated by the active histone mark H3K27ac in another our project which aimed to investigate allelic chromatin interactions in BM. Then, we combined these two different chromatin interaction data with HiFi reads to generate two new BM assemblies. Excitedly, the results demonstrated that most (>98%) of the contigs in these two new assemblies became well-phased without haplotype switches (Fig. 2b, c). More importantly, since the chromatin interaction can be nested across an entire chromosome, when contigs from a single haplotypic assembly were further anchored onto chromosomes, most of them from a same chromosome can be well-phased except for a few regions that overrepresented by short contigs (Fig. 2e). At the same time, the genome redundancy for the assemblies produced by HiFi reads alone has been alleviated in these HiFi + HiChIP assemblies without significant compromise of key assembly metrics (Table 1). We also tested the trio-binning method by using the whole genome short reads from B73 and Mo17 to partition and assemble BM HiFi reads, resulting in near-perfect phased assemblies in which the contigs in two partitions came from B73 and Mo17, respectively (Fig. 2d). As mentioned earlier, the trio-binning strategy is only applicable when the short reads of two parents are available, which cannot be satisfied for the large number of naturally existing plant hybrids. Consequently, we proceeded our subsequent analysis with the two chromosome-scale haplotypic assemblies produced by HiFi and HiC.

### Haplotypic assemblies can resolve the complex loci with parental structural variations

We compared the two haplotypic assemblies with their B73 and Mo17 reference genomes respectively, showing overall good collinearity at chromosomal scales (Fig. 3a-b). However, further analysis is still needed to validate the fine-scale sequence accuracy, especially for loci with extensive structural variations that may interfere the assembly and phasing processes. Therefore, we selected several loci including *ub3*, *gt1* and *ZmSh1-1* which were reported to have extensive SVs in both up and downstream regions between B73 and Mo17 [4, 14], where the sequence of two haplotypic assemblies exhibited ultra-high identity and collinearity with the sequences of B73 and Mo17 (Fig. 3a-c). For loci with more complex copy number variations, the two haplotypic assemblies can even resolve the 27-kDa γ zein locus which has two gene copies in Mo17 resulted from a 15-kb tandem duplication as compared with only one copy in B73 (Fig. 3d). Thereby, we may conclude that most of the complex loci with extensive parental SVs can be properly resolved in our BM haplotypic assemblies.

**Fig. 3.**
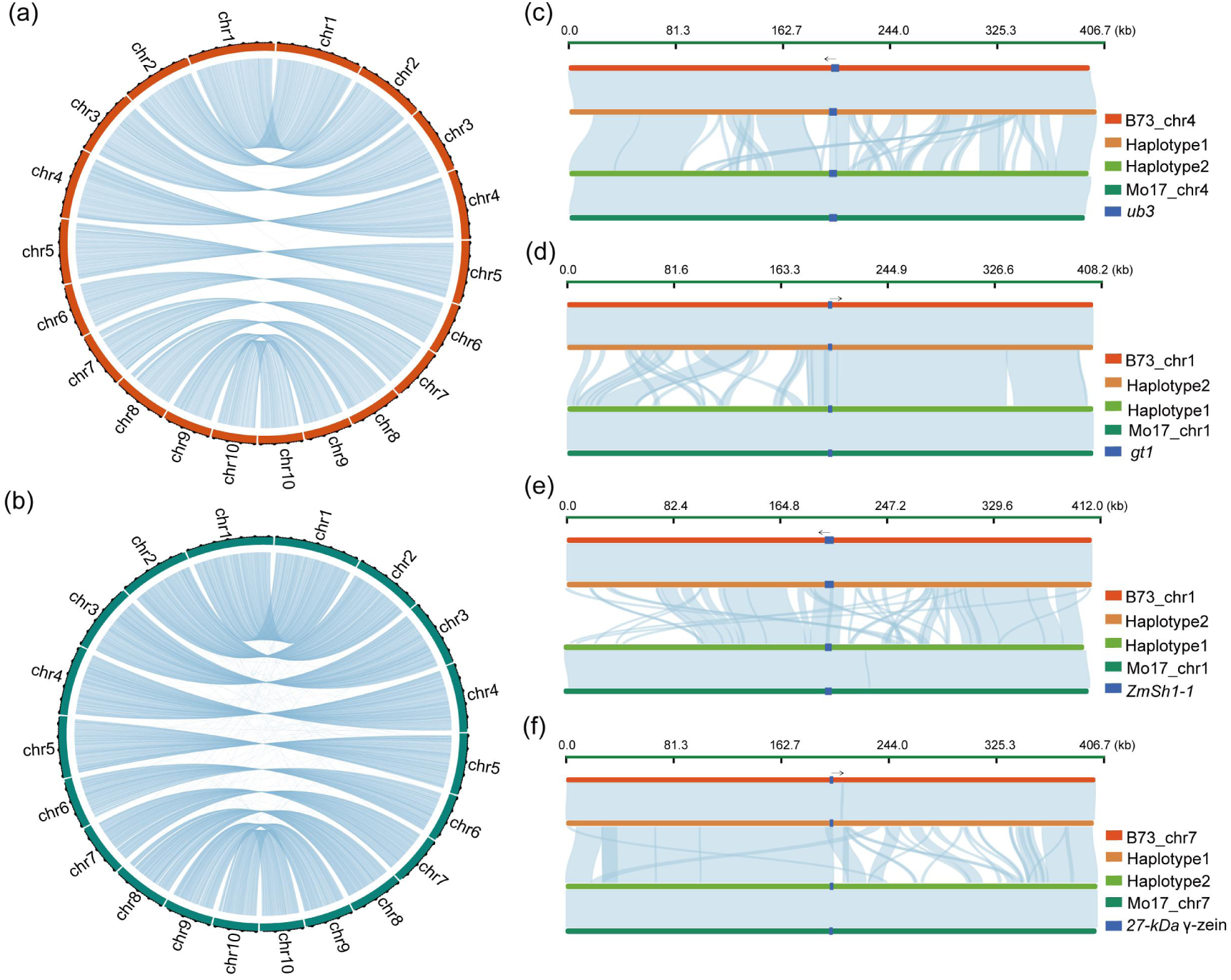
Haplotypic maize assemblies can accurately resolve the loci with extensive parental structural variations. **(a)** Comparison of the collinearity between the ten chromosomes assembled using the HiFi+HiC strategy from the B73 source and the ten chromosomes assembled using the trio-binning strategy. **(b)** Comparison of the collinearity between the ten chromosomes from the Mo17 source and the ten chromosomes assembled using the trio-binning strategy. (c)-(f) For each loci, we exhibited 4 different tracks representing the sequences of B73, two haplotypes, and Mo17. The homologous sequences between two haplotypes are shown as blue bands. The 4 loci are *ub3* (c), *gt1* (d), *ZmSh1-1* (e), and 27-kDa γ-zein (f), respectively.

Our new haplotypic assemblies were expecting to outperform current maize reference genomes in terms of sequence completeness (Table S4 and Fig. S5a-b), therefore a substantial number of sequence gaps may be patched although the newest B73 genome was near finishing (only 151 gaps remained). According to the criteria that a sequence gap can be traversed by a intact contig in our haplotypic assemblies, we found more than half of the gaps (n = 83) can be patched with their length ranged from one kilobases to several hundred of kilobases (Fig. S5a-b). This could be attributed to the improved length, quality and coverage of our latest HiFi reads, and more importantly the well-designed Hifiasm algorithm which was effective in assembling some of the ‘tough’ regions. In terms of the sequence properties, approximately three-quarter of the filled sequences were annotated as repeat elements, with *Copia* retrotransposons contributed predominantly to them (Table S5). Notably, we found a considerable proportion of the filled sequences came from *CentC* elements (Fig. S5c), which was consistent with the evidence from a series of recent gapless assemblies that *CentC* and other complex tandem repeats had been frequently found in the gap filling sequences [23]. In summary, the above results indicated the high accuracy of haplotypic assemblies from the combination of advanced long reads and assemblers, which at the same time can patch many residual gaps in released plant genome assemblies.

### The HiFi+HiC strategy is also efficient for decoding interspecific plant hybrids

Interspecifc hybrids are also common in plants that have played critical roles in their specification, domestication and breeding applications. They usually exhibit higher heterozygsity, along with more extensive parental structural variations over intraspecifc hybrids, raising the necessitity to further verify this HiFi+Hi-C strategy in more complex plant hybrids. Our colleagues manually generated a hybrid by crossing maize B73 with a wild teosinte Ames_21814 (hereafter denoted as BT, heterozygosity estimation of ∼7.7%) and sequenced it by PacBio HiFi and Hi-C technologies. They further applied the Trio-binning method to generate an independent, highly continuous genome assembly of teosinte (∼2,436 Mb & contig N50 of ∼62.3 Mb). By assuming that the parents of BT were unvailable, we choose to directly assemble these HiFi and Hi-C reads and further anchored the haplotypic contigs onto two chromosomal-scale assemblies. Rejoicingly, the two haplotypic assemblies were both well-phased at chromosomal scale as BM (Fig. 4a), showing outstanding sequence quality (BUSCO: 98.2% vs 98.3%; LAI: 32.8 vs 33.0) and maintaining high collinearity with the assemblies produced by Trio-binning method (Fig. S6). These assemblies were also consistent with the trio-binning results that more than 100 Mb novel sequences were found in the teosinte haplotype over B73 haplotype (Fig. 4b), suggesting post-domestication genome contraction or expansion (or introgression) happened in maize and teosintes, respectively. For example, two ultra-large blocks have been identified that specifically existed in teosinte chr2 and chr5 as compared with their homologous B73 chromosomes (Fig. 4c). Altogether, these results suggested that the HiFi+HiC strategy was applicable for generating chromosomal-scale, phased haplotypic assemblies with a varying degree of genome heterozygosity.

**Fig. 4.**
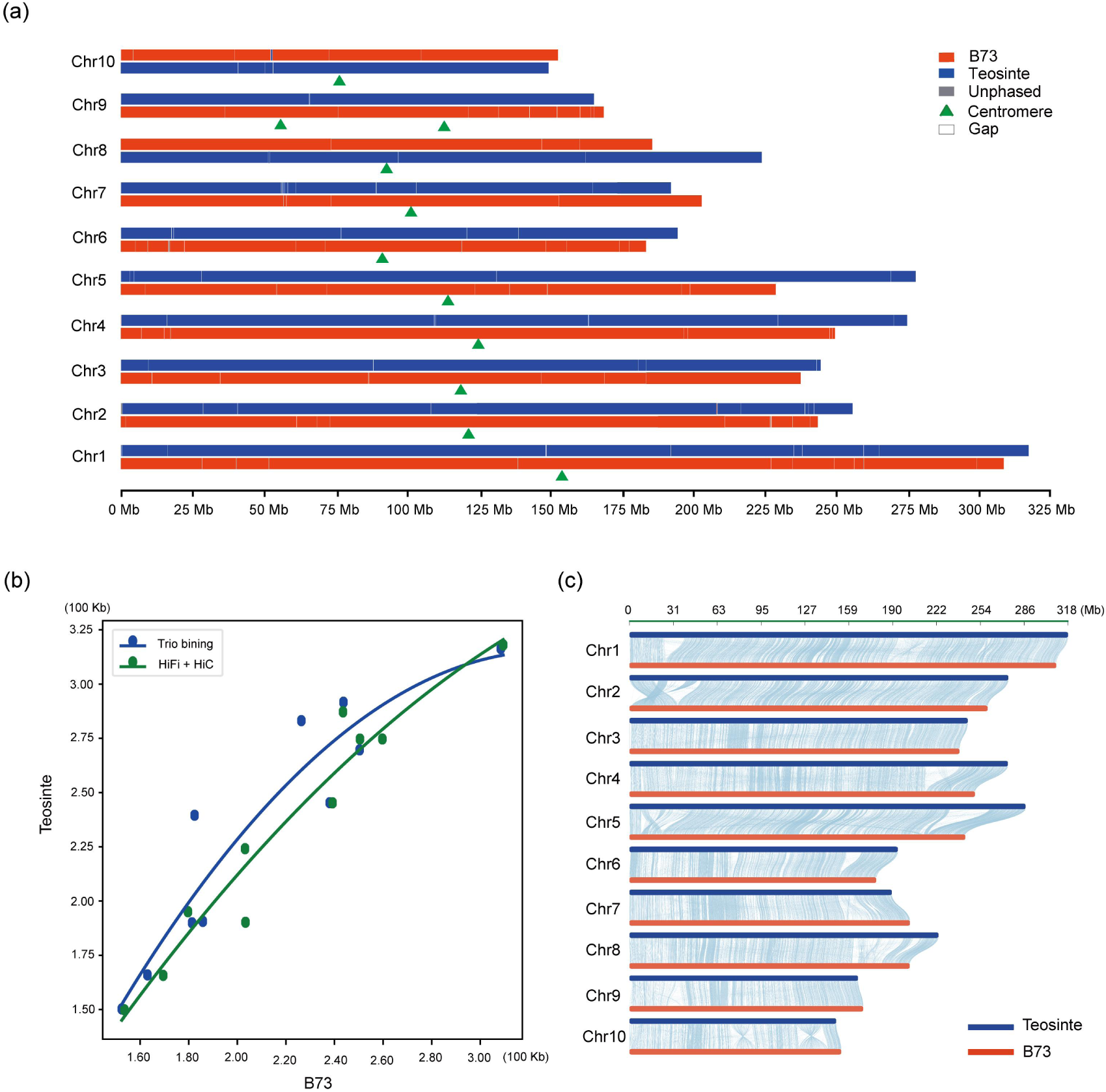
Good performance of the BT assemblies using HiFi+HiC strategy. **(a)** Chromosome-level phasing results for interspecies hybrid assemblies between b73 and teosinte with HiFi and Hi-C. For each chromosome, the top track and the bottom track indicate haplotype 1 chr and haplotype 2 chr, respectively. **(b)** The size of the 10 chromosomes in B73 and teosinte genomes, respectively. **(c)** Comparison of the collinearity between B73 and teosinte genomes.

### Bi-parental graph strategy to represent and analyze haplotypic genomes

Although the haplotypic assemblies generated by HiFi+Hi-C were highly continuous, accurate and well-phased at chromosomal scales, they cannot be directly used as genome reference for downstream analysis until their sequence redundancy and allelic variants can be properly processed. Graph genome has been recently demonstrated as an important strategy to incorporate populational genome sequences for improved reads mapping, structural variation calling, genome-wide association studies (GWAS) and many other downstream applications [7]. We here integrated our BM haplotypic assemblies into a genome graph, which we called the bi-parental graph, by using the haplotype1 as backbone and the SNPs, small InDels (< 50 bp) and other structural variations (SVs, >= 50 bp) between two haplotypes as alternative edges. In terms of small variations, a total of 5,428,920 SNPs and 720,086 small InDels between two haplotypes have been identified by MUMmer [27]. Although the coordinates between hap1 backbone sequences and B73 genome cannot be unified for cross-validation, the number of SNPs and InDels and their chromosome distribution were highly similar between hap1-hap2 and B73-Mo17 comparisons (Table S6), suggesting overall good representation of these parental SNPs and InDels. As for large SVs, they were indenpendently called by cuteSV [28], SVIM-asm [29], Sniffles [30] and Assemblytics [31], then those supported by at least two callers were retained (n = 63,205). Finally, a bi-parental genome graph incorporating hap1 sequences and all high-confidence genetic variants was contructed by VG [32], and at the same time an simplified graph that specifically designed for mapping RNA-seq reads was contructed by HISAT2.

We first accessed the performance of this graph by aliging the resequencing short reads from B73, Mo17, BM, and other maize inbred lines (PH207, OH43 and Chang7-2) to the graph using VG, and compared their alignment rates with the commonly used aligner BWA MEM [33] using both hap1 and hap2 linear sequences as reference. Generally, VG achieved an enhanced alignment rate (∼26% to ∼32%) compared with those aligned by BWA (mapping quality > 0, Fig. 5a). The BWA alighment rate was especially dropped for OH43 and PH207, two vital germplasm that had far genetic distance with B73 and Mo17 [34]. By contrast, the alignment rate by VG remained steady across the 6 short-reads datasets (Fig. 5a). We also aligned the RNA-seq reads from BM 12-day embryo to hap1, hap2 linear references and the bi-parental genome graph using HISAT2, finding significantly increased mapping accuracy since reads without sequence mismatches were significantly elevated (92.2%, Fig. 5b) by using genome graph as compared with those of two linear references (83.9% and 83.0%).

**Fig. 5.**
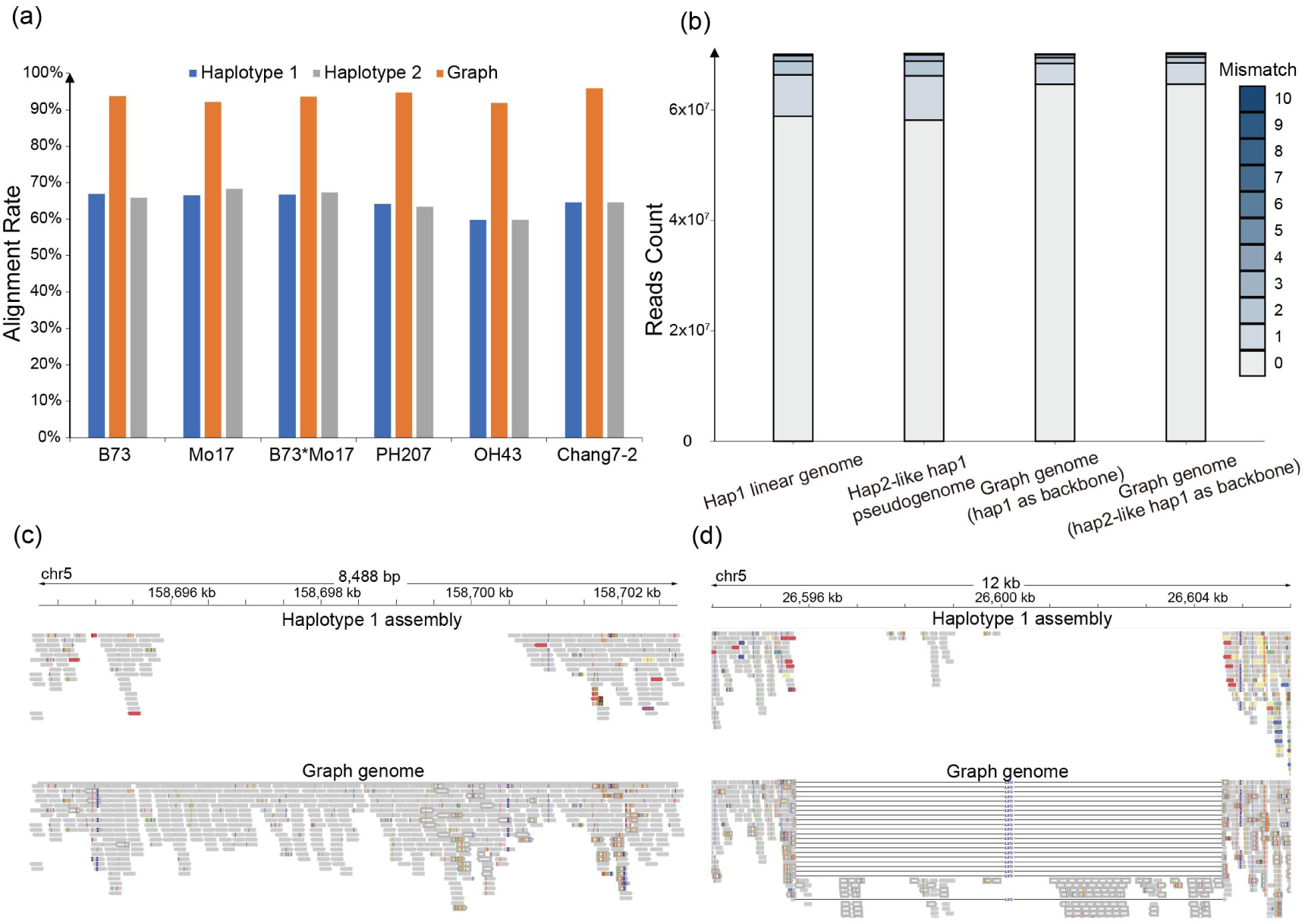
Graph genome improved alignment rate and accuracy, and enhanced capability of genotyping complex SVs. **(a)** The whole genome short sequences of B73, Mo17, BM hybrid, PH207, OH43 and Chang7-2 were aligned to our haplotype 1 assembly and graph genome, respectively, to compare the alignment rate of the two references. **(b)** The mismatch numbers in the situation that BM hybrid RNA-seq were mapped to the linear and graph references. **(c)** One example that more PH207 short DNA reads can be mapped to the corresponding region when graph genome was used as reference. Upper and bottom represent the mapping detail of the situations when linear and graph genome were aligned to, respectively. **(d)** One example that more PH207 short DNA reads can be mapped to the corresponding region when graph genome was used as reference. Upper and bottom represent the mapping detail of the situations when linear and graph genome were aligned to, respectively. Structural variations between haplotype 1 and haplotype 2 assembly existed in this region, so short reads exhibited totally different alignment situations. Black lines denote another alternate path (deletion) of graph genome.

Since bi-parental graph was more superior in aliging short reads than linear genome, we expect it to have improved sensitivity and accuracy for SV identification. As a demonstration, when short reads of PH207 were aligned to the hap1 linear reference by BWA, we found a valley of read depth that spanning 8.5 kb, suggesting potential absence of this region in PH207. However, when aligning to the bi-parental graph using VG, this region can be covered by enough reads with excessive of SNPs and small Indels (Fig. 5c), clarifying the existence of a homologous sequence which differed substantially with hap1 and more closely resembled hap2. Another case was the identification of a large deletion (∼8.9kb) in PH207 with many “jumping” reads when aligned againt graph, by contrast no reads can “jump” across this deletion when aligned to hap1 linear reference (Fig. 5d). These results demonstrated the outperformace of bi-parental graph as new reference for downstream genome analysis, for example SV calling, using cost-effective short reads.

### Bi-parental genome graph was applicable for analyzing allele-specific gene expression

Allele-specific expressed genes are fundamentally important in deciphering heterosis and imprinting, although their identification remained still challenging to date. When RNA-seq reads from a hybrid were mapped onto a linear reference genome, reads from one parent that more genetically distant with the reference may suffer more mapping bias, then introduce false positives for the identification of allele-specific expressed (ASE) genes. A general approach to alleviate these mapping bias was to substitute the reference genome with the genotypes of parental SNPs to construct a “pseudo-genome”, which was conceptually similar to the graph genome approach and has been demonstrated to be highly efficient [35]. Inspired by the pseudo-genome strategy, we have devised an innovative approach called ‘pseudo-graph genome’ to combine the advantages of pseudo-genome and graph genome. In this approach, the haplotype2-like pseudo-haplotype 1 with replaced genotypes was served as the backbone to construct the graph, whose alternate paths were integrated by the genotypes at SNP loci from haplotype 1 and the indels between haplotype 1 and haplotype 2.

To verify the superiority of ‘pseudo-graph genome’, we compared the performance of two linear genomes and two graph genomes (Fig. 6a) in identifying ASE genes and assessing mapping bias. These included the haplotype 1 linear genome, the haplotype 2-like haplotype 1 pseudo-genome, the graph genome (haplotype 1 as the backbone), and the graph genome (haplotype 2-like haplotype 1 as the backbone). Initially, we applied HISAT2 [36], a versatile aligner to map the RNA- seq reads of BM embryo to the above four reference genomes respectively, and quantified the number of reads originating from haplotype 1 and haplotype 2 sequences by utilizing their shared SNPs within each gene. Under ideal conditions without mapping bias, the rate of reads derived from haplotype 1 should approach approximately 50%. The results revealed that, superior than two linear genomes, the graph genome, especially the pseudo-graph genome, attained a nearly perfect mapping balance at a global level (Fig. 6b). Besides, we identified allele-specific SNPs under our rigorous identification criteria (Method). Remarkably, when linear genome served as reference, the allele-specific SNPs displayed strong bias towards either haplotype 1 or haplotype 2 sequences (Fig. 6c). In contrast, using the graph could identify a much more comparable number of allele-specific SNPs from two haplotypes (Fig. 6c). Then, we further compared their performances under varying levels of stringency in ASE gene identification criteria. It showed that, irrespective of the stringency, the pseudo-graph genome consistently outperformed the two linear genomes showing minor mapping bias (Fig. S8). Moreover, most of the ASE genes identified by graph genome were supported by linear genomes (Fig. 6d), highlighting potentially high false positives in linear genomes. To sum up, the graph genome, particularly the pseudo-graph genome, boasts high mapping accuracy (Fig. 5b) and low mapping bias (Fig. 6b, c and Fig. S8), proving it emerges as a robust reference alternative to linear genome.

**Fig. 6.**
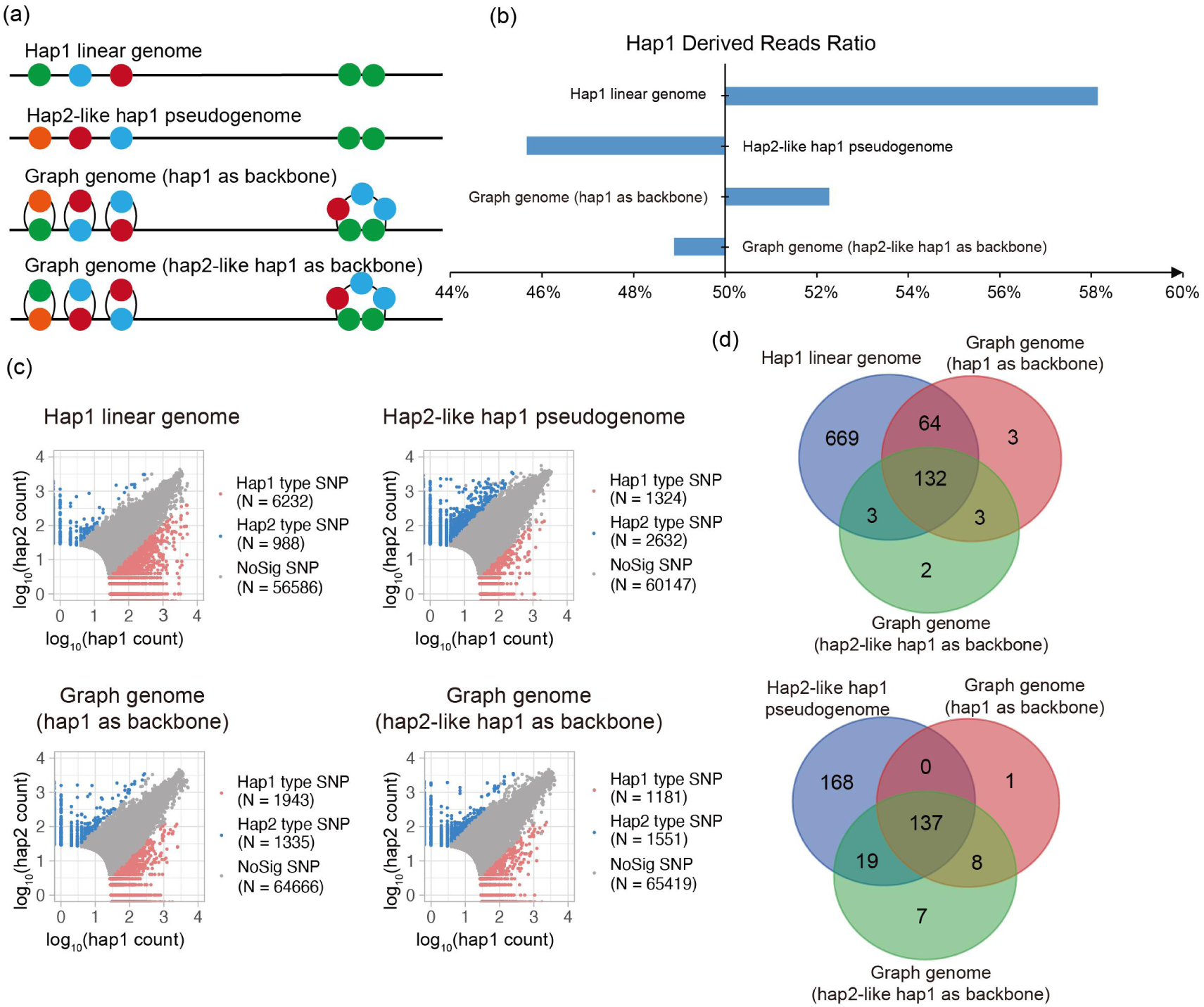
Allele-specific expressed genes identified when linear genome and graph genome used as reference, respectively. **(a)** A schematic diagram showing the four different reference genomes. **(b)** The ratio of hap1 derived reads (Methods) to total reads at analyzable SNPs in the circumstances of four different references. **(c)** The comparison of the SNP calling bias among four assembling strategies. Each dot denotes an analyzable SNP with 30 more informative reads. The x-axis and y-axis are the number of hap1-derived reads and hap2-derived reads at each SNPs taking the log 10. Gray dot (Balanced) denotes SNP without showing preference for hap1 or hap2 genotype. Red dot (Hap1 biased SNP) denotes SNP showing preference for hap1 genotype, at which site hap1 derived reads takes up more than 70% of the total reads. Same thing goes with the blue dot (Hap2 biased SNP). **(d)** The venn plots of the allele-specific-expressed genes called from different assembling strategies. On the left and right are the outcome overlap of the three methods used to identify hap1 and hap2 allele-specific-expressed gene, respectively.

### Summary & Discussion

Decoding the genomes of hybrids has long been a tough work despite of its great significance, as a substantial number of naturally existing hybrids are waiting to be assembled. Although some pioneer efforts have been made to assemble a couple of hybrid plant genomes, they may still lack proper phasing and in-depth analysis, preventing the development of a unified approach to fully harness the potential of these hybrid genomes. Therefore, despite the trio-binning strategy commonly used with known parential assemblies, we combined the Hifi and HiC data to generate the chromosome-scale, well-phased assemblies of highly heterozygous maize genome under circumstance where parential assembilies were unnecessary. In addtion, we proved that the HiFi+HiC strategy could resolve the complex loci with parental structural variations and work effectively for decoding interspecies hybrid genome, facilitating the analysis of wild plant genetic resources. Furthermore, we upgratded the hybrid graph genome for maize with the HiFi+HiC data and carried out the ‘pseudo-graphgenome’ to boasts high mapping accuracy and low mapping bias, which shed light into the analysis of allele-specific expressed genes in terms of hybrid assemblies. However, although our analysis showed clear advantages in identifying candidate ASE genes, we cannot further classify their parent-of-origins since those identified as haplotype 1 types included both B73 and Mo17 specifically expressed genes. To solve this issue, the upcoming well-phased hybrid genomes will hold more promise to be applied in the identification of ASE genes and the complex imprinting gene studies.

As progress made in our study, we still found that haplotype switches were inevitable when sequencing reads, even though the up-to-date long HiFi reads, were assembled directly without the participation of other data. However, we demonstrated that the integration of these two switched haplotypic assemblies into a genome graph might be a good solution that can enable more efficient variant calling, allele-specific expression interference and many other potential analyses. In the future, important progress should be taken to use haplotype sensitive long-distance interactions, for example the Hi-C data, to alleviate the haplotype switch rate and ultimately phase the two haplotypes at genome-wide scales. With confidence, we anticipate the graph genome approach may play an important role in utilizing these hybrid assemblies to understand heterosis, along with the surge of assembling hybrid plant genomes in the coming years.

## Materials and Methods

### Plant material

The maize (*Zea mays*) hybrid BM (B73 * Mo17) used in this study was grown in a greenhouse at 25 ℃ in dark conditions for 14 days, and the above-ground portions of seedlings were harvested and frozen in liquid nitrogen immediately for genomic DNA extraction. High molecular weight genomic DNA was extracted from isolated nuclei for library construction.

### Genome sequencing

Genomic DNA was sequenced by the Pacific Biosciences (PacBio) Sequel platform at the Frasergen company, Wuhan, China. Six SMRTbell libraries were constructed and sequenced by the PacBio Sequel II system, yielding a total of 80 Gbp raw reads, corresponding to ∼60X genome coverage.

### Assembly and evaluation

Approximately 5 million PacBio SMRT reads of maize hybrid were assembled and tested by different parameters (-s) of Hifiasm (v0.19.5) [12]. The assembly generated with default parameters was retained due to presenting the highest contiguity. BUSCO (v5.4.2) [25]was further used to evaluate the genome-assembly completeness. ‘embryophyta_odb10’, which contained 1,614 single-copy orthologous genes was used as a searching dataset. The BUSCO was run by Python3.8.5 following the code of ‘busco3.0.2/bin/run_busco −m geno −i genome.fa −o out_name −l embryophyta_odb9 −c 15’. The LAI was calculated following the pipeline of LTR_retriever (v2.9.0) (https://github.com/oushujun/LTR_retriever). Mo17 and B73 genomes were assessed through the same method.

### Haplotype switches identification

Contigs in the genomes larger than 100Kb were phased by Merqury (v1.3) based on the parental specific k-mers and were classifiedaccording to the proportion of hap-mer in total number (>= 0.7). At the same time, the two haplotypes were compared to B73 and Mo17 genomes with Minimap2 (v2.24), respectively, and the unphased contig (< 0.7) were further classified according to coverage and consistency.

### Analysis of repetitive elements and gap filling

RepeatMasker (https://github.com/rmhubley/RepeatMasker) was used to discover and identify repeats in the hybrid genome with the curated maize TE libraries derived from the Maize TE Consortium (MTEC). The situations of gaps in the B73 and Mo17 reference genomes were filled with the primary or alternate contigs as follows: primary or alternate contigs were aligned to the genomes of B73 and Mo17 by MUMmer (v4.0.0) [27] with the parameter ‘nucmer −l 100 −c 500’. In addition, if the primary or alternate contigs can map to the flanking region of a gap in B73 or Mo17, then they were considered to be the situation of filling this gap.

### Identification of variations

We identified SNPs and Indels polymorphisms (indels, length < 50 bp) between haplotype 1 and haplotype 2 assembly with MUMmer (v4.0.0) with the parameter ‘nucmer −l 100 −c 500’. Then, SVs (length > 50bp) were detected by Assemblytics (http://assemblytics.com/) [31]. To get more accurate SVs information between haplotype 1 and haplotype 2 assembly, we also mapped the alternate contigs sequence to the primary pseudochromosome with minimap2 using the parameters ‘minimap2 - ax asm5’. A total of four callers: Sniffles (v2.0.7) [30] with the parameters ‘sniffles -- minsvlen 50 −t 50 --mapq 20 --minsupport 1 --long-del-length 100000 --long-dup-length 100000’, SVIM-asm (v1.4.2) [29] with the parameters ‘svim-asm haploid -- min_mapq 20 --min_sv_size 50’ and CuteSV (v2.24) [28] with the parameters ‘cuteSV −t 70 --min_mapq 20 --min_size 50 --max_split_parts −1 --max_size 100000 --min_support 1’ were used for variant calling. Only SVs supported by at least two callers and where the callers agreed regarding the type of variant were retained for downstream analysis.

We also conducted several strategies to eliminate false discovery of SNPs. Nearby SNPs were removed using sliding window method: if a 10-bp width, 1-bp step window contains >= 3 SNPs, then we removed the SNPs and INDELs in them. Additionally, we mapped the hybrid PacBio HiFi reads to the haplotype 1 assembly using Minimap2 (v2.17), and the BAM file generated was filtered with the mapping quality threshold of 30. The SNP we identified before would be discarded if it was covered with less than 3 heterozyfous reads.

### Graph genome construction

The variation graph toolkit (vg) pipeline [32] was used for the construction of graph genome, with the information including SNPs, indels and SVs between the two assemblies identified previously. The vg pipeline was also used for variant mapping with short reads to evaluate the performance of the graph genome, comparing the linear primary assembly using the standard BWA-MEM2 (v2.2.1) pipe [33].

### RNA-seq data processing

The embryo RNA-seq data (Gene Expression Omnibus (GEO) database accession no. GSE95399) [37] of the B73 * Mo17 hybrid was used to perform transcriptome mapping comparison between the linear and graph genome. Raw reads were firstly filtered by fastp (v0.22.0) with default parameters. The filtered high quality reads were aligned to both the linear and graph genome with HISAT2 (v2.2.1) [36]. PCR duplicates were removed with Samtools (v1.3.1).

### Allele-specific expression analysis

The ASE gene identification pipeline was displayed in Fig. S7. Firstly, the HISAT2’s out bam file was transformed to pileup format with Samtools (v1.3.1) to get the detailed mapping information at the SNP sites between the haplotype 1 and haplotype 2 assembly. Thus, genes whose exon didn’t contain SNP sites were not analyzable. Hap1 derived reads and hap2 derived reads at each SNP site were summed up respectively. For SNPs covered by thirty more informative reads (reads whose mapping cigar don’t have ‘.,atcgATCG’ but others were excluded), Chi-square test was performed to assess the deviation from an expected hap1:hap2 ratio of 1. Additionally, to account for the many SNPs being evaluated simultaneously and independently, *p*-values are also corrected to reduce the false discovery rate using the Benjamini-Hochberg method. SNPs were filtered under the following criterias: (1) the ratio of preferred genotype derived reads to the total reads were more than 0.85; (2) adjusted *p*-values under 0.01 were considered significantly hap1 or hap2 biased. If a gene’s exon regions contain 3 more the same biased SNPs in the same direction, then it was identified as a ASE gene. The code and scripts for ASE analysis are available at Github (https://github.com/yjiang296/ASE-gene-classify).

## Acknowledgements

We express our gratitude to Xingguo Wu from Professor Yongrui Wu’s lab from the Shanghai Key Laboratory of Plant Molecular Sciences for their valuable assistance and helpful disscusion in the assembly of the BT genome.

## Author contributions

J.S. designed this research project. Q.K. and Y.J. analyzed most of the data and prepared figures. X.G., Z.W. and Y.L. participated in the data analysis and interpretation. Q.K., ZH.W. and Y.G. involved in the data collection. Q.K. and Y.J. wrote the manuscript. J.S. and ZJ.W. finalized the manuscript. All the authors have read and approved the manuscript.

## Data availability statement

The PacBio HiFi data has also been deposited in the NCBI (https://www.ncbi.nlm.nih.gov) with accession number SRR22252681, SRR22252683, SRR22342922, SRR22269434 and SRR22269435. All sequencing data for creation of the teosinte line haplotype were downloaded at the NGDC (https://ngdc.cncb.ac.cn) under the accession number SAMC874385, SAMC873392, SAMC874386 and SAMC874387.

## Supporting information

**Fig. S1** The distribution of read length (top) and GC content (bottom right) of the 118 Gb PacBio HiFi reads generated in this study.

**Fig. S2** Evaluation of the genome assembly quality among different sequencing depths of PacBio HiFi data.

**Fig. S3** Comparison of the genome size and contig N50 of two haplotypic assemblies under different −s threshold in Hifiasm.

**Fig. S4** Examples of phasing regions between the haplotype 1 and haplotype 2 contigs.

**Fig. S5** The B73 gaps filled by two haplotypic assemblies.

**Fig. S6** Two haplotypic assemblies were both well-phased at chromosomal scale and having high collinearity with the assemblies produced by Trio-binning method.

**Fig. S7** The workflow of the allele-specific-expressed gene identification in this study.

**Fig. S8** Allele-specific-expressed gene bias under different identification thresholds.

**Table S1** Heterozygosity estimation by k-mer analysis of maize and other hybrids.

**Table S2** Summary of sequencing data of the BM genome.

**Table S3** Comparison of genome assembly statistics.

**Table S4** The sequence gaps in B73 that have been filled in two assemblies.

**Table S5** The composition of transposable element in filled-gaps.

**Table S6** The number of SNPs, InDels and SVs between haplotype 1-haoplotype 2 and B73-Mo17.

## Supplementary Figures

**Fig. S1.**
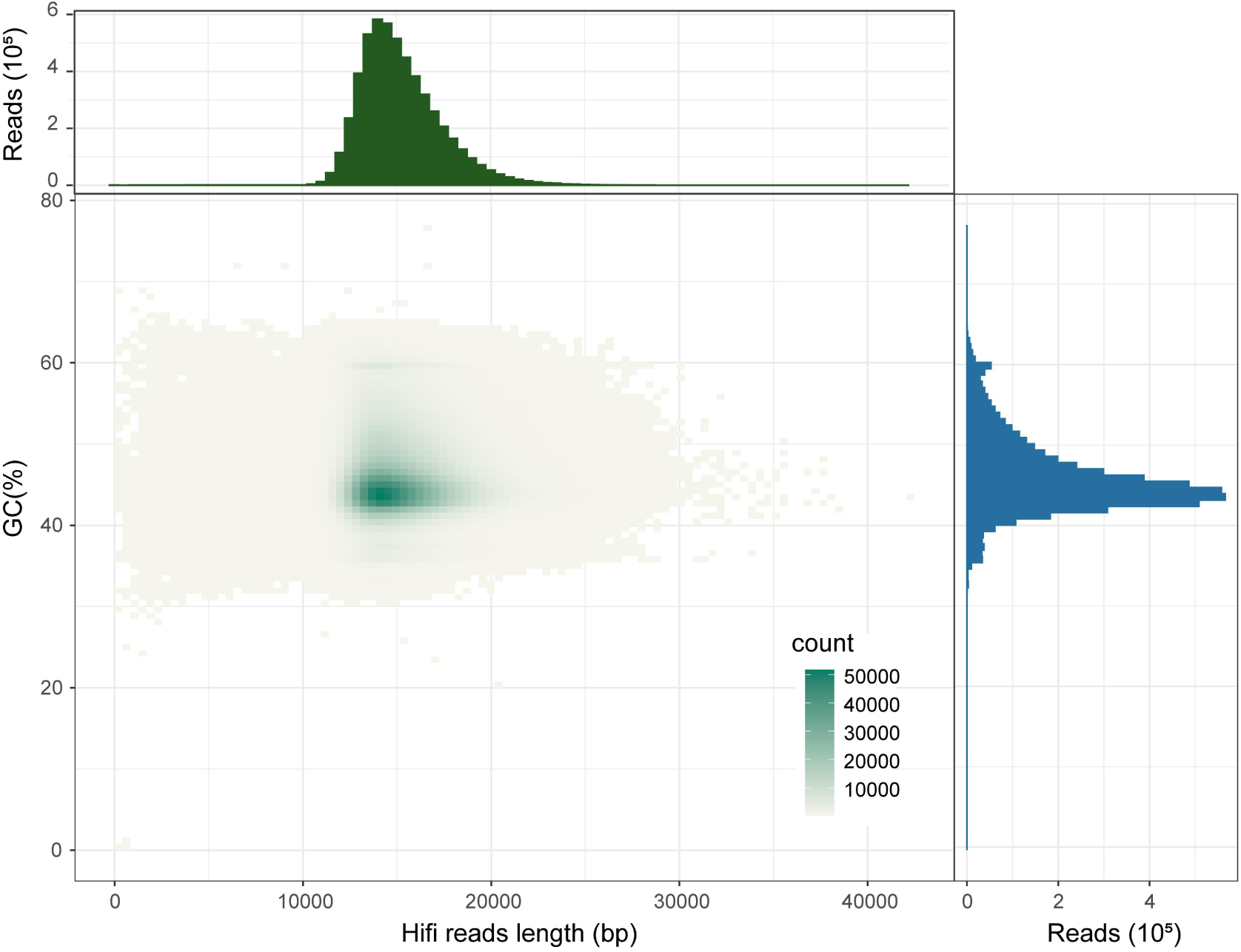
The distribution of read length (top) and GC content (bottom right) of the 118 Gb PacBio HiFi reads generated in this study.

**Fig. S2.**
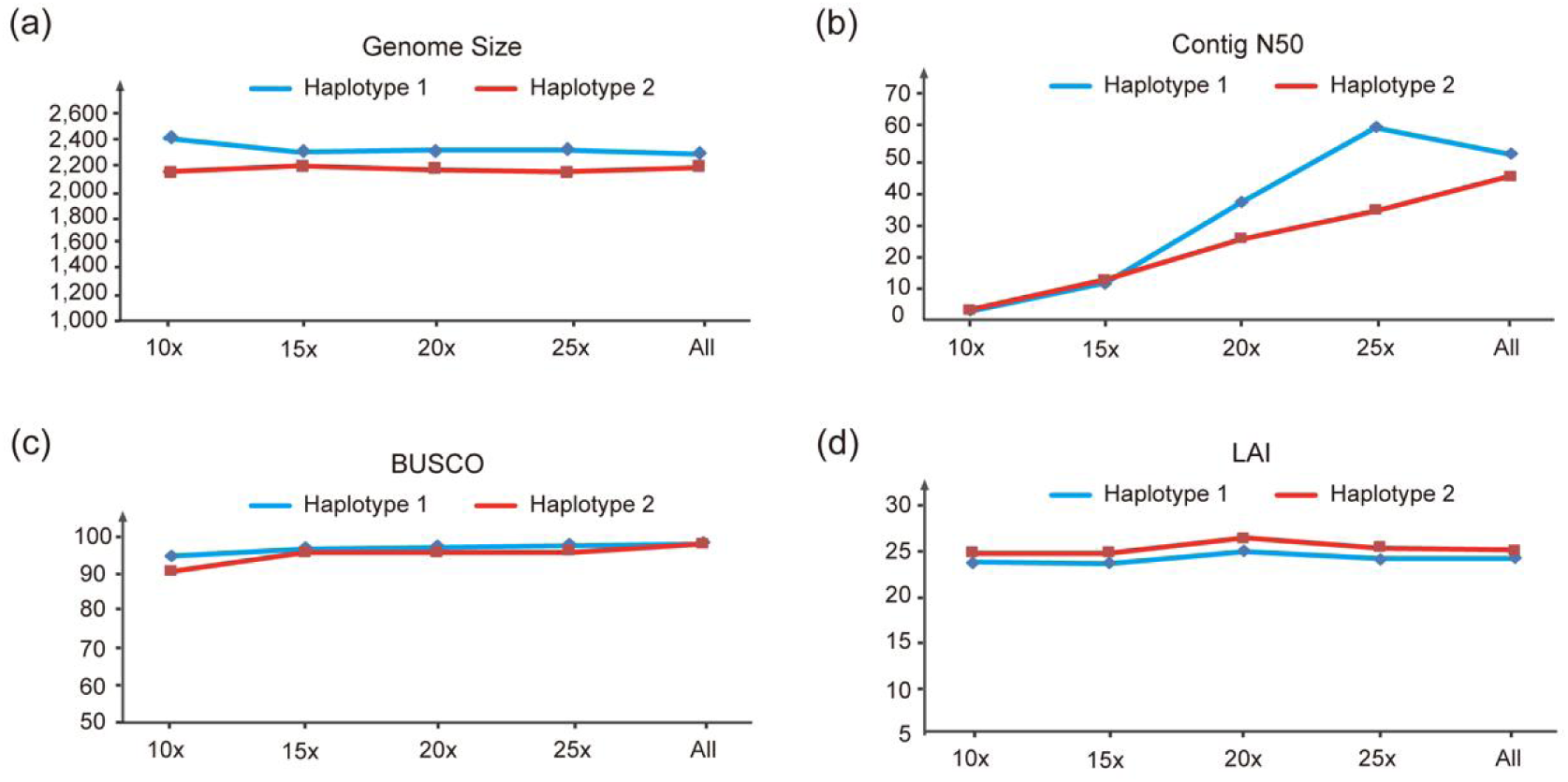
Evaluation of the genome assembly quality among different sequencing depths of PacBio HiFi data. **(a)** The genome size and **(b)** contig N50 of two haplotypic assemblies. The x-axis represents the depth of PacBio HiFi data. The y-axis represents the length (Mb) of genome sequencing of each haplotype. **(c)** The BUSCO and **(d)** LAI of two haplotypic assemblies.

**Fig. S3.**
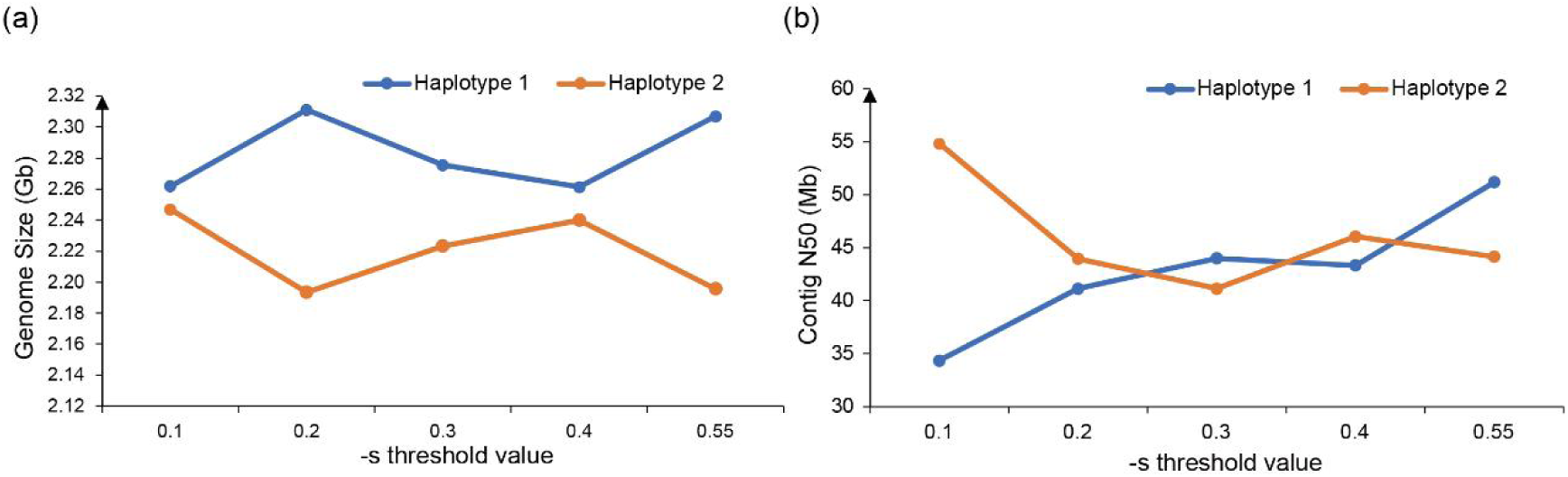
Comparison of the genome size and contig N50 of two haplotypic assemblies under different −s threshold in Hifiasm.

**Fig. S4.**
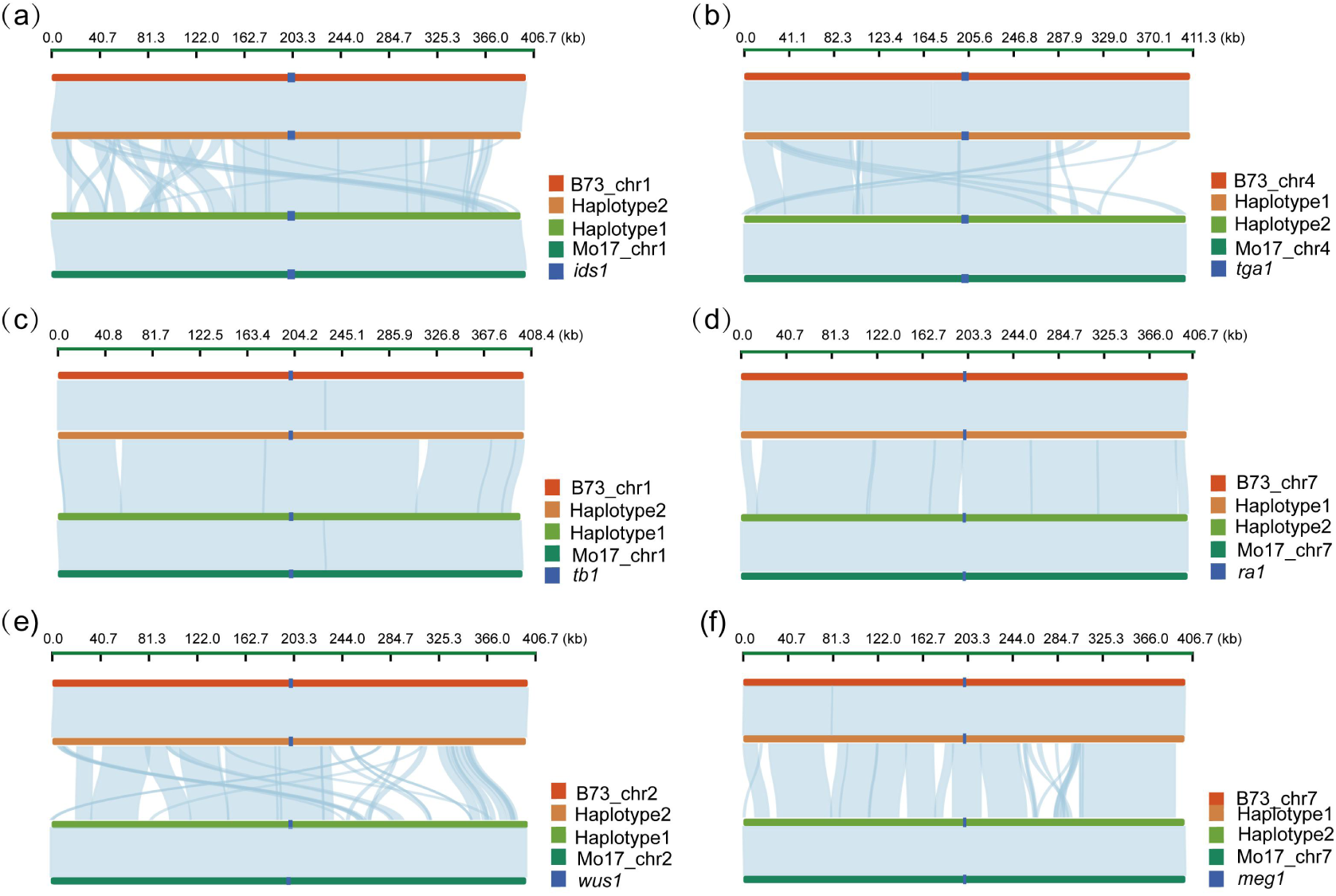
Examples of phasing regions between the haplotype 1 and haplotype 2 contigs. Collinearity between primary and alternate assembly in the 200kb region upstream and downstream of **(a)** *ids1*, **(b)** *tga1*, **(c)** *tb1*, **(d)** *ra1,* **(e)** *wus1*, **(f)** *meg1* and collinearity between two haplotypes and parent sequences. Red blocks (topside) represent B73 genome, while bottle-green blocks (bottom) are the Mo17 genome. The middle blocks are primary and alternate and the blue blocks are gene sequence.

**Fig. S5.**
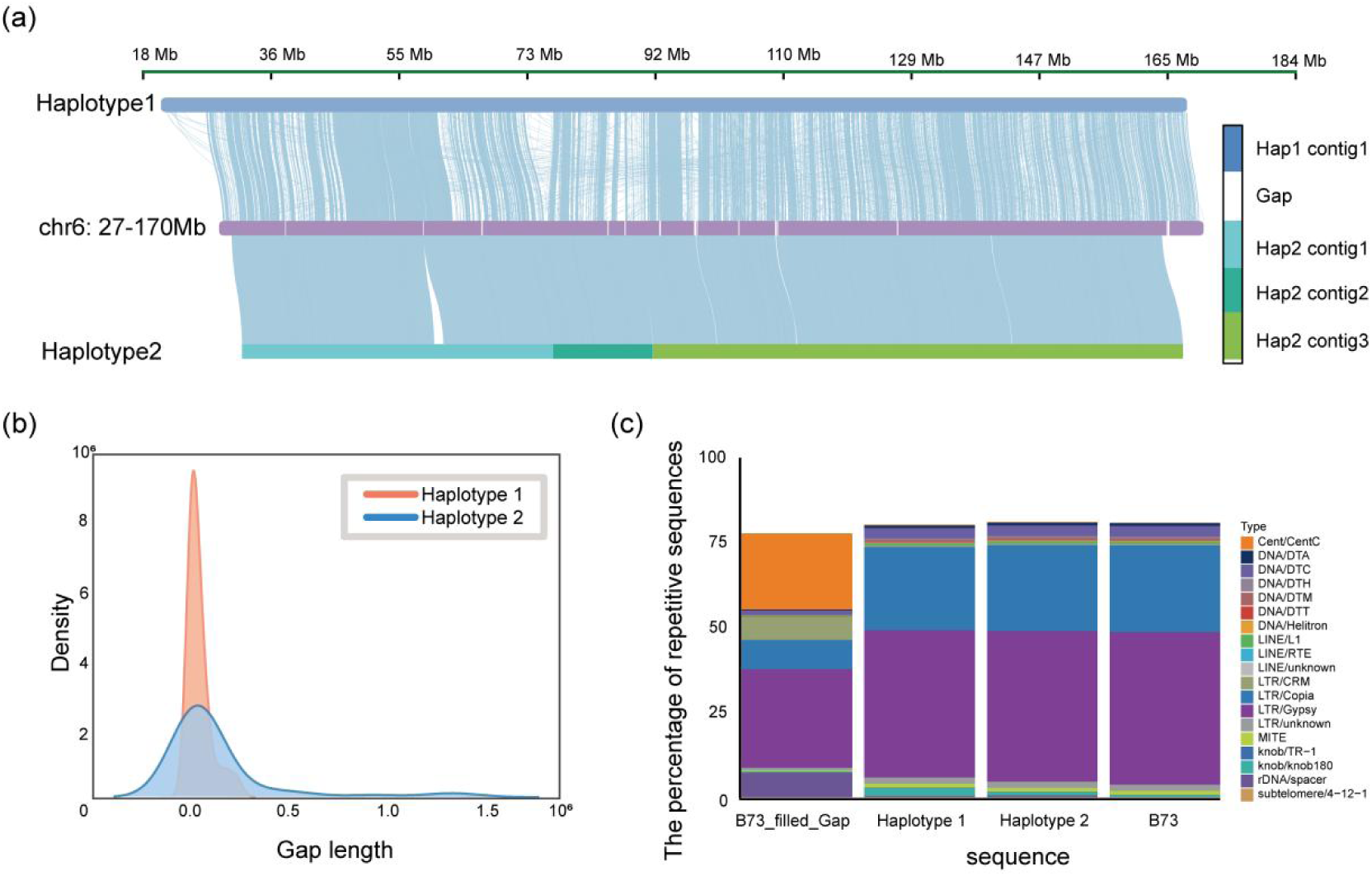
The B73 gaps filled by two haplotypic assemblies. **(a)** Local sequence filling of **the** B73 reference genomes. **(b)** The length distribution of filled gaps from B73. **(c)** The composition of transposable elements in filled-gaps sequences.

**Fig. S6.**
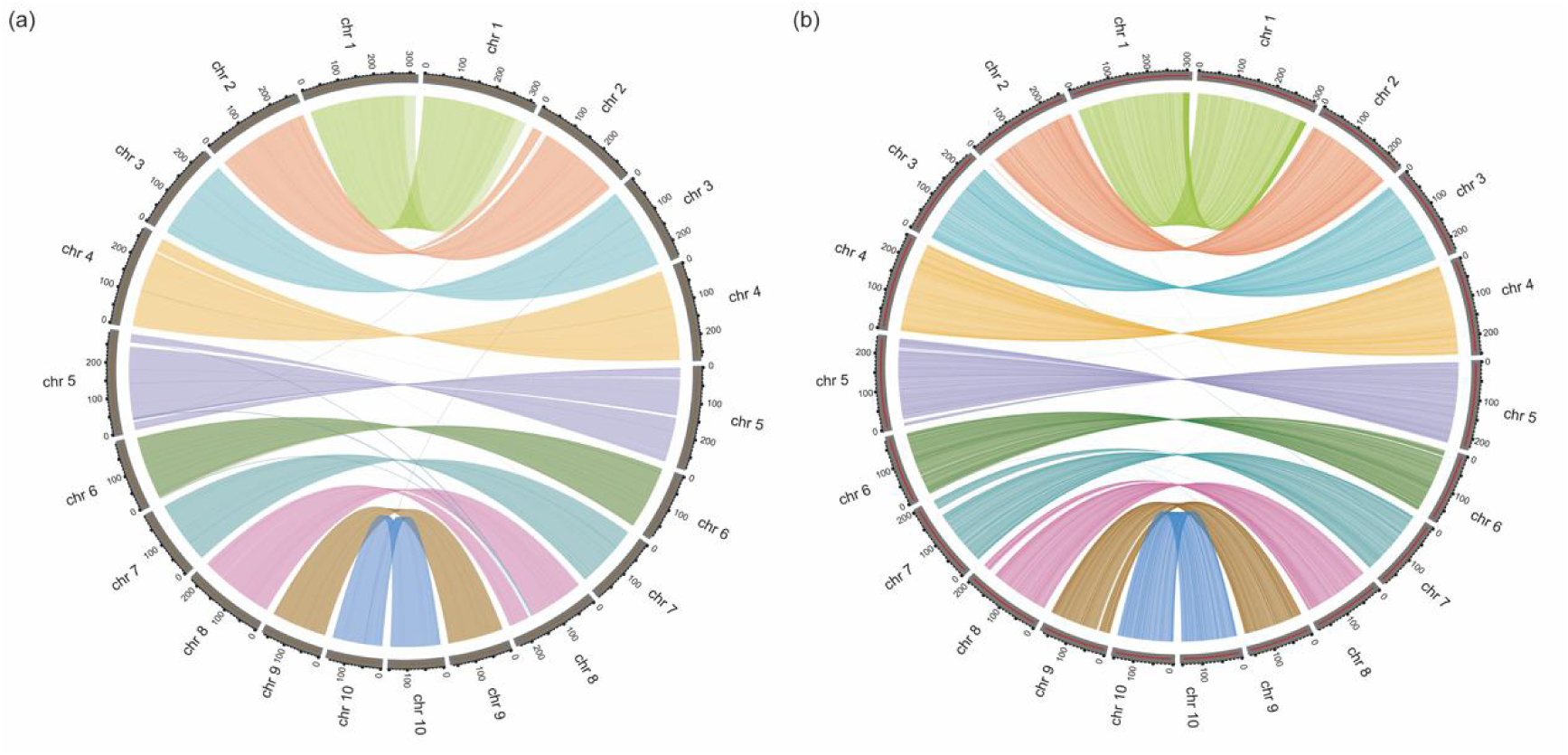
Two haplotypic assemblies were both well-phased at chromosomal scale and having high collinearity with the assemblies produced by Trio-binning method. **(a)** Comparison of the collinearity between the ten chromosomes assembled using the HiFi+HiC strategy from the B73 source and the ten chromosomes assembled using the trio-binning strategy. **(b)** Comparison of the collinearity between the ten chromosomes from the teosinte source and the ten chromosomes assembled using the trio-binning strategy.

**Fig. S7.**
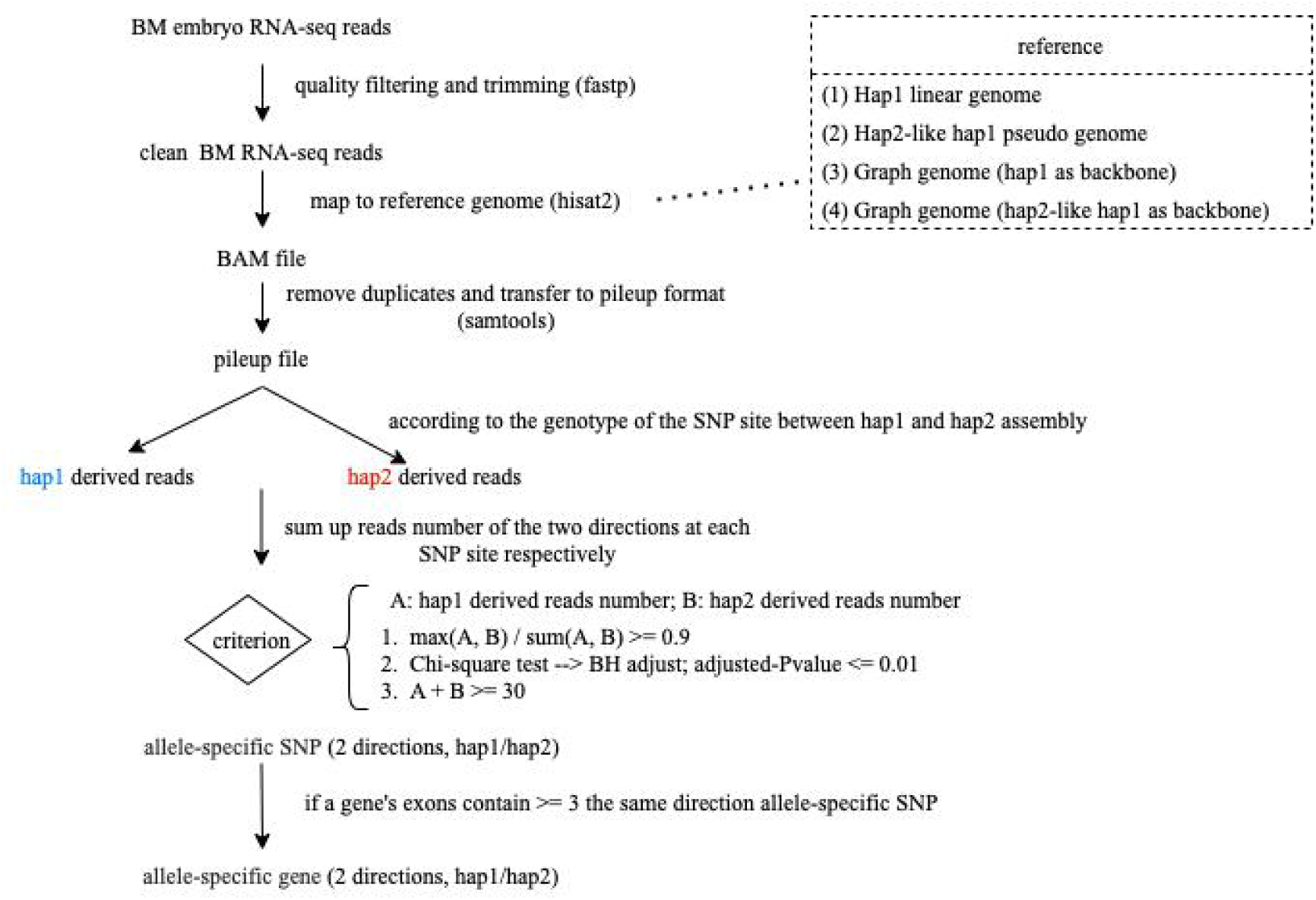
The workflow of the allele-specific-expressed gene identification in this study.

**Fig. S8.**
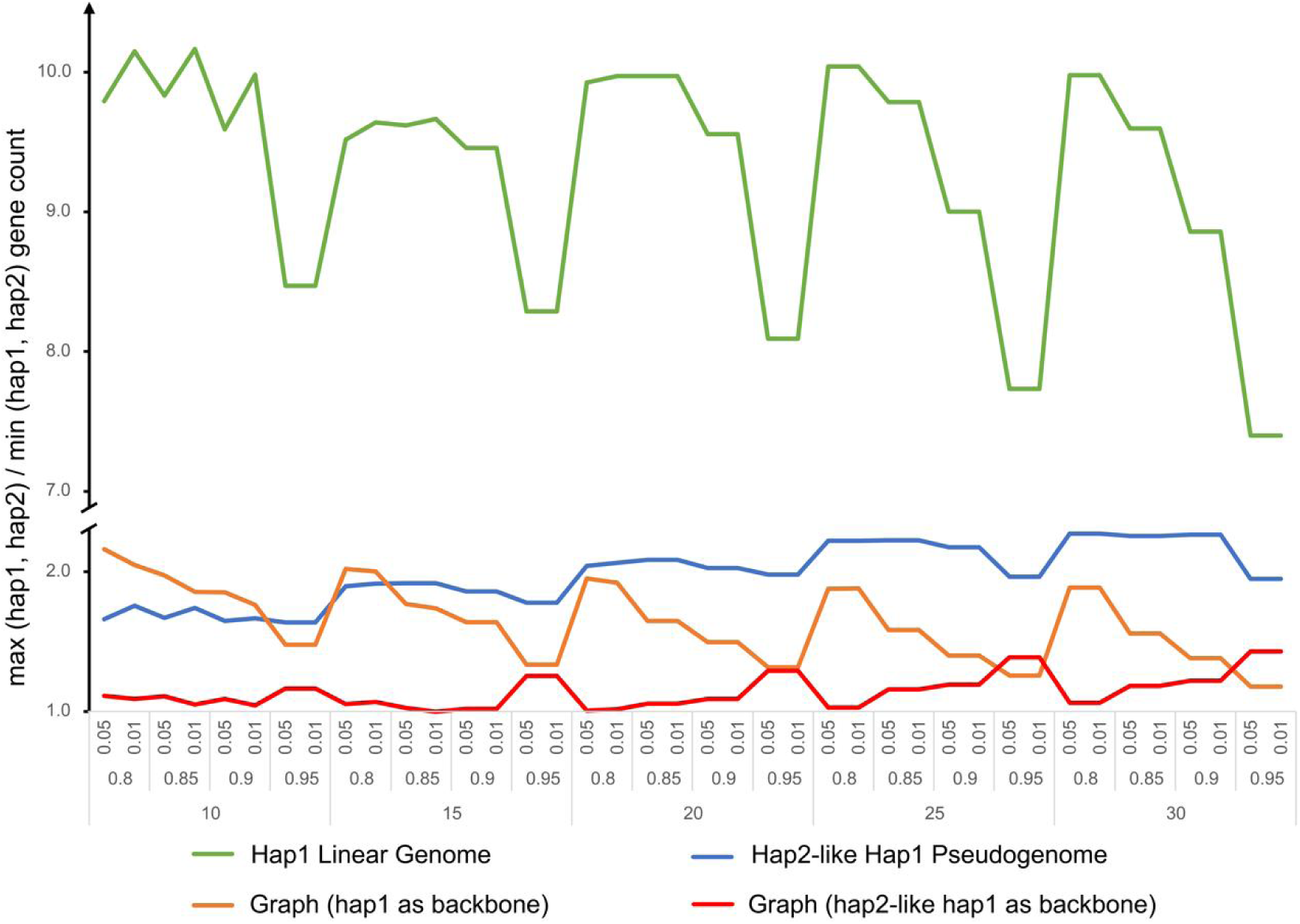
Allele-specific-expressed gene bias under different identification thresholds. X-axis represents the groups of allele-specific SNP identification threshold: (1) at least how many informative reads for an analyzable SNP (15, 20, 25 or 30); (2) the ratio of preferred genotype derived reads to the total reads (0.8, 0.85, 0.9 or 0.95); (3) adjusted *p*-value (0.05 or 0.01). Y-axis represents the value of max (hap1-allele-specific genes count, hap2-allele-specific genes count) / min (hap1-allele-specific genes count, hap2-allele-specific genes count).

## Supplementary Tables

**Table S1.**
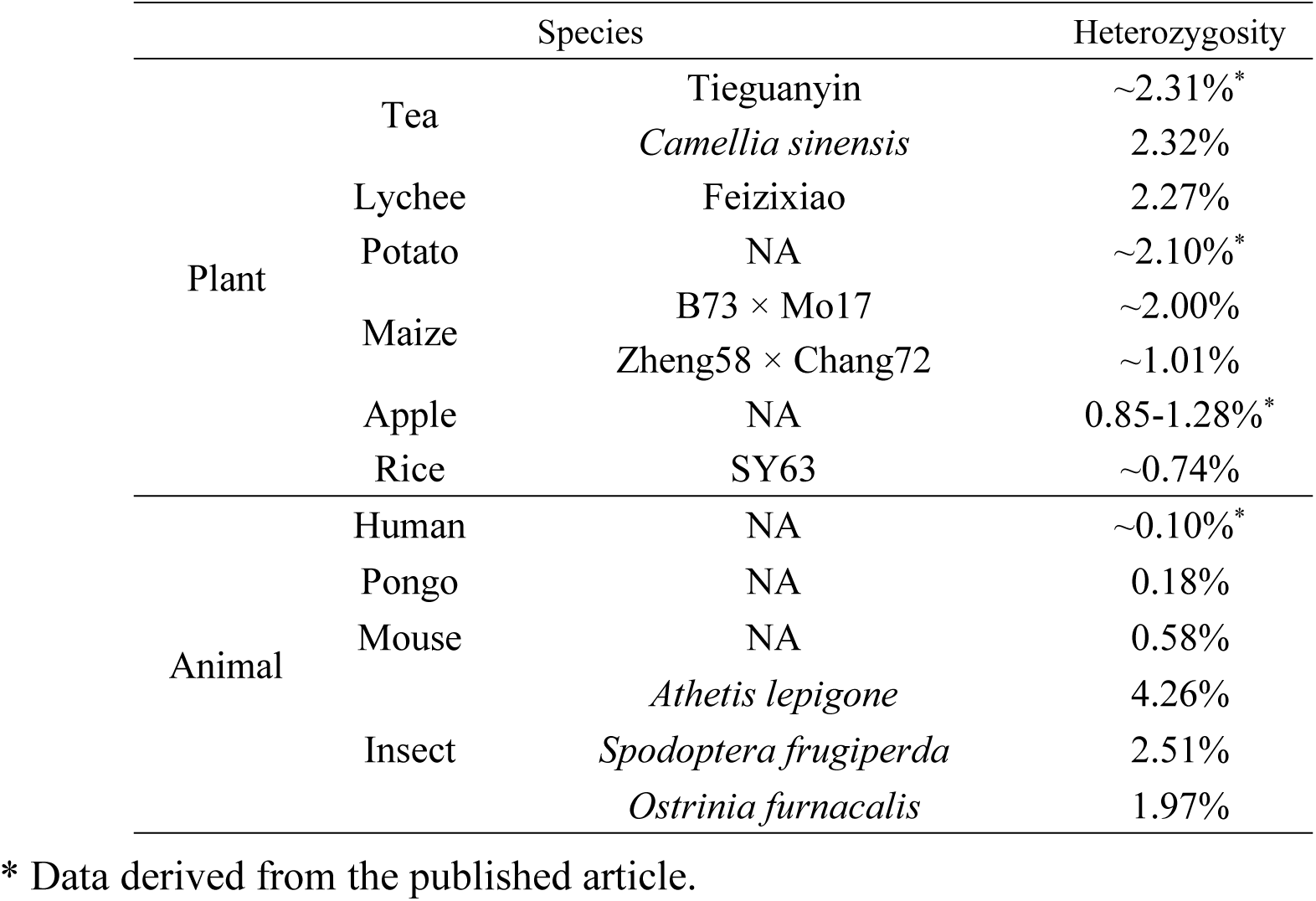
Heterozygosity estimation by k-mer analysis of maize and other hybrids.

**Table S2.** Summary of sequencing data of the BM genome.

| **Dataset** | **Flow cell number** | **Reads Number** | **Total length (bp)** | **Genome depth\*** | **N50 length of reads (bp)** |
| --- | --- | --- | --- | --- | --- |
| **PacBio HiFi data** | 6 | 7,368,408 | 118,026,484,198 | 54.2 | 16,090 |
| **Hi-C data** | - | 642,683,864 | 96,269,576,394 | 44.2 | 150 |
| **Hi-Chip data** | - | 466,419,256 | 69,962,888,400 | 32.1 | 150 |

\* Calculated with the B73 reference genome size of 2,178 Mb.

**Table S3.**
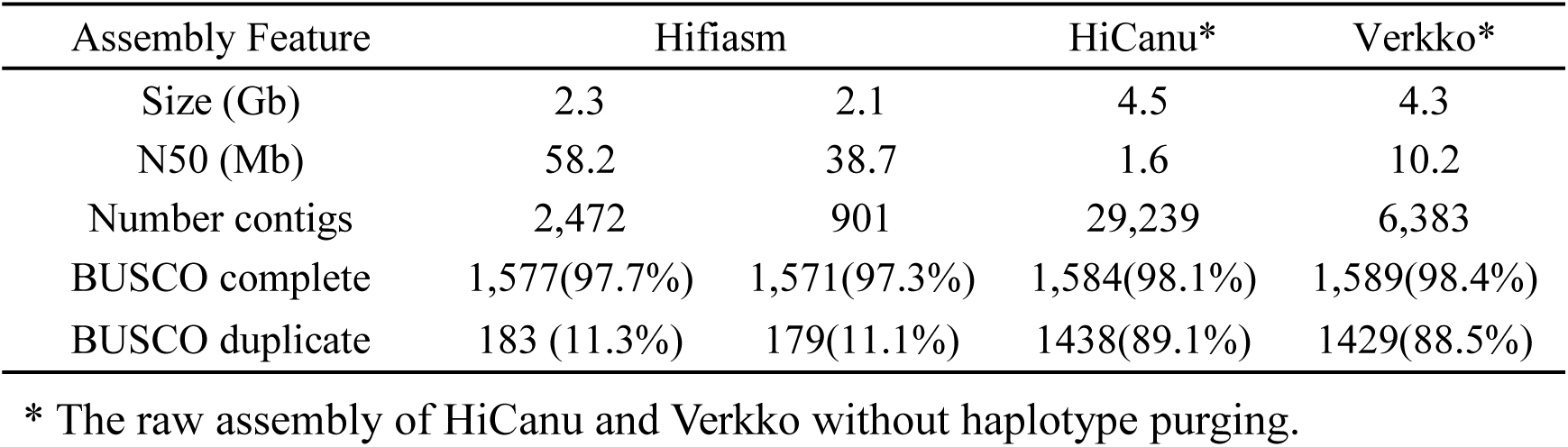
Comparison of genome assembly statistics.

**Table S4.**
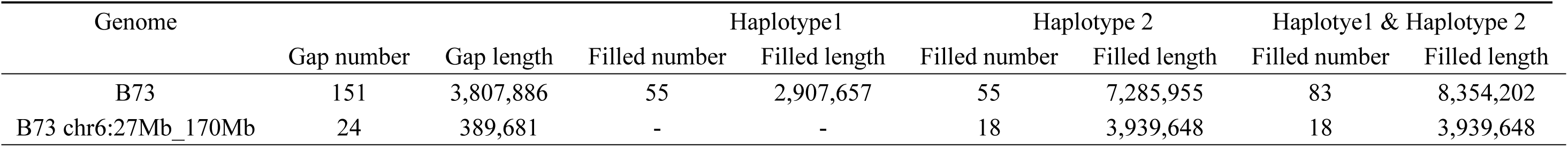
The sequence gaps in B73 that have been filled in two assemblies.

**Table S5.** The composition of transposable element in filled-gaps.

| Classification | Length (bp) | Proportion (%) |
| --- | --- | --- |
| Cent/CentC | 1,873,968 | 22.43 |
| DNA/DTA | 21,414 | 0.26 |
| DNA/DTC | 108,924 | 1.30 |
| DNA/DTH | 4,134 | 0.05 |
| DNA/DTM | 13,031 | 0.16 |
| DNA/DTT | 0 | 0.00 |
| DNA/Helitron | 6,705 | 0.08 |
| LINE/L1 | 23,002 | 0.28 |
| LINE/RTE | 0 | 0.00 |
| LINE/unknown | 7,918 | 0.09 |
| LTR/CRM | 557,617 | 6.67 |
| LTR/Copia | 721,810 | 8.64 |
| LTR/Gypsy | 2,420,546 | 28.97 |
| LTR/unknown | 59,091 | 0.71 |
| MITE/DTA | 13,831 | 0.17 |
| MITE/DTC | 1,459 | 0.02 |
| MITE/DTH | 8,242 | 0.10 |
| MITE/DTM | 108 | 0.00 |
| MITE/DTT | 1,313 | 0.02 |
| knob/TR-1 | 0 | 0.00 |
| knob/knob180 | 34,002 | 0.41 |
| rDNA/spacer | 601,341 | 7.20 |
| subtelomere/4-12-1 | 0 | 0.00 |
| Total | 6,478,456 | 77.55 |
| Filled gaps length | 8,354,202 | - |

**Table S6.** The number of SNPs, InDels and SVs between haplotype 1-haoplotype 2 and B73-Mo17.

| Assembly | SNPs | InDels | SVs |
| --- | --- | --- | --- |
| Haplotype 1 vs Haplotype 2 | 5,428,920 | 720,086 | 20,984 |
| B73 vs Mo17 | 5,508,854 | 721,852 | 20,431 |

